# Nanoparticle endocytosis is driven by monocyte phenotype rather than nanoparticle size under high shear flow conditions

**DOI:** 10.1101/2023.06.29.547038

**Authors:** Dasia A. Aldarondo, Chris Huynh, Leah Dickey, Colette Bilynsky, Yerim Lee, Elizabeth C. Wayne

## Abstract

Monocytes are members of the mononuclear phagocyte system involved in pathogen clearance and nanoparticle pharmacokinetics. Monocytes play a critical role in the development and progression of cardiovascular disease and, recently, in SARS-CoV-2 pathogenesis. While studies have investigated the effect of nanoparticle modulation on monocyte uptake, their capacity for nanoparticle clearance is poorly studied. In this study, we investigated the impact of *ACE2* deficiency, frequently observed in individuals with cardiovascular complications, on monocyte nanoparticle endocytosis. Moreover, we investigated nanoparticle uptake as a function of nanoparticle size, physiological shear stress, and monocyte phenotype. Our Design of Experiment (DOE) analysis found that the THP-1 *ACE2*^-^ cells showed a greater preference for 100nm particles under atherosclerotic conditions than THP-1 wild-type cells. Observing how nanoparticles can modulate monocytes in the context of disease can inform precision dosing.

## Introduction

Atherosclerosis is a chronic inflammatory vascular condition characterized by subendothelial plaque formation caused by the accumulation of fats, cholesterol, and cells^1^. Subendothelial accumulation of oxidized low-density lipoproteins (LDLs) upregulates endothelial cell expression of monocyte recruitment and adhesion membrane receptors^2,3^. Once monocytes infiltrate the plaques, they differentiate into macrophages. These macrophages oxidize LDLs and cholesterol-forming foam cells that augment the inflammatory signaling^4^. Macrophages continue to uptake the cholesterol, which is esterified and stored as lipid droplets in the endoplasmic reticulum, eventually becoming engorged and undergoing endothelial reticulum stress, apoptosis, or necroptosis^3,5^. Ultimately, these vulnerable plaques rupture, leading to other cardiovascular diseases such as thrombosis and stroke^3^.

Monocytes and macrophages are members of the mononuclear phagocyte system (MPS), which accounts for a significant portion of nanoparticle clearance^6,7^. Endocytosis of nanoparticles by monocytes depends upon various factors including nanoparticle morphology (size and shape) and monocyte phenotype.^8,9^ The size and shape of nanoparticles impact the surface interactions that facilitate exposure of the monocyte with the nanoparticle carrier composition or cargo. In contrast, monocyte phenotype determines the surface receptors and stimuli which will dictate interactions^10,11^.

Cellular endocytosis is classified into two large umbrellas, phagocytosis or pinocytosis dependent on the uptake mechanism ^6,8,12^. Large particle sizes ranging from 250 nm to 3 μm are endocytosed via a phagocytosis^12,13^. Pinocytosis is a broader classification with many sub-classifications determined by the receptor interactions.^12,14^. The main groups of pinocytosis are micropinocytosis, Clathrin-dependent, caveolin-dependent, and clathrin and caveolin-independent endocytosis^12^. Macro-pinocytosis is the largest classification of pinocytosis, taking up particles ranging in size from 3nm -200nm^9^.

Macro-pinocytosis internalizes particles through extensions of the plasma membrane^15^. Mid-range particles are internalized using clathrin-mediated (50nm – 150nm) or caveolin receptor-mediated endocytosis (50nm – 250nm)^16^.

Monocyte phenotype can be described through cytokine and receptor expression. Patients afflicted by disease typically have altered phenotypic populations^17^. Monocyte phenotypes exist on a broad spectrum but are usually classified into three overarching classifications based on CD16 and CD14 expression. The three groups are classical (CD14+/CD16-), Intermediate (CD14+/CD16+), and non-classical (CD14dim/CD16++). Classical monocytes comprise most of the monocyte populations, while intermediate and non-classical combined comprise a small population.

In Atherosclerosis, an increase in intermediate and non-classical indicates Atherosclerosis progression^18,19^. Increased frequency of non-classical monocyte is associated with better cardiovascular outcomes, while increased classical and intermediate frequency is associated with worse cardiovascular outcomes^20–22^. Many adhesion receptors also play a significant role in Atherosclerosis, such as CCR2, CXCR1, and CCR5, which monocyte use to bind to their respective ligand on the endothelium for plaque progression^23,24^. Many other cytokines are implicated in the development of Atherosclerosis. While there is significant research on the role of nanoparticle size on endocytosis in the immortalized human monocytic THP-1 cells, these dynamics are less studied in disease-impaired monocytes.

Another critical enzyme and receptor implicated in Atherosclerosis is Angiotensin-converting enzyme 2 (ACE2). ACE2 is a crucial mediator in the Renin-angiotensin signaling pathway, maintaining homeostasis within the cardiovascular system^25,26^. ACE2 primarily converts Ang-II into Angiotensin 1-7 (Ang1-7), a vasodilating and antiproliferative peptide hormone^27^. Ang (1-7) works to counter the inflammatory and proliferative effects caused by Ang-II, produced by the Angiotensin-converting enzyme (ACE) ^27^. ACE2 is present throughout cardiovascular tissues, including smooth muscle cells, endothelial cells, lung epithelial cells, and monocytes. Loss of *ACE2* expression is correlated with hypertension^28,29^, and increased heart failure^30^. In cardiovascular diseases (CVDs), ACE expression on the monocyte surface is upregulated and contributes to a pro-inflammatory phenotype^31^.

Many atherosclerosis studies utilize nanoparticles to encapsulate existing therapeutics or block receptors important to monocyte plaque infiltration^32^. Still, these nanoparticles are often systemically delivered, leading to systemic effects that may impact functions essential to other cellular processes elsewhere in the body, causing various side effects^33^. Despite the advancement in research, there has been limited translational success in nanotherapeutics^34^. In *in vitro* research, there is significant research on the role of nanoparticle size on endocytosis in the immortalized human monocytic THP-1 cells, these dynamics are less studied in disease-impaired monocytes. This study aimed to investigate how altering *in vitro* monocyte models impacted the endocytosis of different-size nanoparticles. To approach this, we utilized an *ACE2* knockdown THP-1 cell line to mimic a diseased cell phenotype, as well as exposed cells to physiological shear stresses to study the uptake of polystyrene nanoparticles.

## Materials and Methods Materials

### Materials

THP-1 monocytes were purchased from ATCC. The *ACE-2* shRNA lentivirus used was purchased from Santa Cruz Biotechnology; (sc-41400-V). For qPCR analysis all DNA primers were created and purchased from Integrated DNA Technologies, Inc. RNA extraction kits were purchased from Qiagen, and both the High-Capacity cDNA Reverse Transcription system and the PowerTrack SYBR Green Master Mix kit were purchased from Applied Biosystems. For flow cytometry flow cytometry staining buffer were purchased from Invitrogen, as well as the polystyrene nanoparticles (F8887) used in all studies.

### Cell Lines

A stably transduced THP-1 ACE2 knockdown cell line was generated using an *ACE-2* shRNA lentivirus (THP-1-*ACE2*^*-*^). All cell cultures were maintained in RPMI media supplemented with 10% (v:v) fetal bovine serum at 37 °C, and 5% CO_2_in a humidified incubator. 2 mg/mL of polystyrene nanoparticles were added immediately to 10mL culture aliquots by direct addition and gentle mixing before applying shear.

### Shear Stress Experiments

To apply shear to the cells, we used a TA instrument DHR 2 Rheometer (Waters; Delaware, USA) using a double gap geometry. TRIOS software was used to set up a protocol consisting of a flow and peak holds step for 20 minutes. The instrument was run at room temperature.

Both cell lines were passaged to a concentration of 1×10^6^ cells/mL. Each cell line was sheared at 5 dyne/cm^2^ or 40 dyne/cm^2^ resulting in two sample groups per cell line. For each sample, 10 mL of the respective cell culture spiked with NPs was obtained and sheared. After shearing, the culture was transferred into a flask and incubated at 37 °C and 5% CO_2_ for 4 hours. Samples for RNA expression were taken for nanoparticle sizes 20nm, 100nm, and 200nm, and nanoparticle endocytosis flow samples were taken for 20nm, 100nm, 200nm, and 500nm nanoparticles. Samples were taken before applying the shear, directly after application of the shear, and after the four-hour incubation.

### qPCR Measurements

Quantitative polymerase chain reaction (qPCR) was performed on the cell lysate samples to quantify changes in RNA expression of the following genetic markers: TNF-α, MAC-1, ICAM-1, CCR2, CCL2, CD40, VCAM-1, IL-12, ACE2, NOS2, and IL-10. DNA primers. RNA was extracted from cell samples using the RNeasy mini extraction kit. The extracted RNA was converted to cDNA. Following RNA conversion to cDNA, SYBR green was used per the supplied protocol. qPCR was run on a ViiA 7 real-time PCR system (Life Technologies; California, USA) utilizing the preprogrammed SYBR green protocol in the accompanying software. Fold change was measured using a ΔΔCt analysis with 18s as the reference gene. Samples were run in triplicate and are a culmination of 2-3 individual experiments. Data analysis was completed using Excel.

### Flow Cytometry

Culture samples collected at each time point were removed from the media and washed once with PBS. Samples were then fixed on ice in a 4% formaldehyde fixation solution for 15 minutes. Following fixation, samples were diluted with stain buffer, pelleted, and resuspended in 200 µL stain buffer. Finally, samples were plated in a 96-well plate and run through a NovoCyte® flow cytometer (Agilent; California, USA) to measure nanoparticle uptake by the cells. FCS Express 7 cytometry software (De Novo Software; California, USA) was used to measure % positive cells and develop mean fluorescence intensity.

### Design of Experiment Analysis

We expected four factors to significantly impact the cell’s nanoparticle uptake during shear studies. These four factors include THP-1 monocyte phenotype, nanoparticle size, resting time, and magnitude of shear stress. To pinpoint which of these factors significantly affect this percent of cells positive for nanoparticle uptake, we utilized JMP 17 Statistical Analysis Software (SAS) to organize four classical, two-level main effects screening design of experiments (DOEs). Each of the designs had 12-runs and consisted of identical levels of (i) Cell Type: wildtype vs ACE2 knockout, (ii) Shear stress: 5 vs 40 dyne/cm^2^, and (iii) Resting Time: 0 vs 4 hours. The four designs did, however, vary in the levels selected for (iv) nanoparticle size to exploit the potential impact of size competition between a multitude of diameter size ranges. Nanoparticle size levels selected for each of the four models are as follows: (Model A) 20 vs 100 nm; (Model B) 20 vs 200 nm; (Model C) 100 vs 200 nm; (Model D) 100 vs 500 nm. Raw data values reported are the percentage of cells positive for uptake of 100 (Models A, C, and D) or 200 nm (Model B) nanoparticles (Supplemental Tables 1 and 2).

To improve the root-mean-square error (RMSE) of our initial screening models, we employed an Augmented Design to increase the number of experimental runs from 12 to 36, with three replicates of each run from the original DOEs. Experimental conditions and response values for each of the Augmented Designs are provided in Tables S2 and S3. We used JMP’s design of experiments analytical software to fit a Partial Least Squares Linear Regression (SLSLR) model to our data.

## Results and Discussion

### ACE2^-^ THP-1 cells have an increased percentage of nanoparticle endocytosis nanoparticles in high shear stress

Monocyte nanoparticle endocytosis has previously been correlated with shear stress caused by physiological blood flow conditions^35-36^. In the presence of an atherosclerotic plaque the flow conditions as varied as well as the circulating monocyte populations^36,37^. We wished to investigate whether monocytes deficient in *ACE2* would endocytose nanoparticles differently based on flow regimes. In this section, we measured the effect of shear stress, nanoparticle size, and cell phenotype on monocyte endocytosis.

Wildtype and *ACE2* deficient (*ACE2*^—^) THP-1 monocytes were exposed to nanoparticles under high (40 dyne/cm^2^) or low (5 dyne/cm^2^) shear stress conditions using a double gap rheometer. Using flow cytometry, we analyzed the percentage of monocytes positive for nanoparticles (NP Positive %) and the mean fluorescence intensity (MFI). In addition, we calculated these parameters immediately after shear (t = 0) and after a 4–hour rest period to investigate sustained effects.

At higher shear rates, nanoparticle endocytosis increased significantly in ACE2^-^ cells compared to static conditions at both time points, with an approximate 20-30% increase in the percentage of cells positive **(Figure 1 C-F)**. Interestingly, there was still no observable nanoparticle size dependence on the number of cells that take up particles. After the rest period, significantly more ACE2^-^ cells to up cells in the presence of 20nm particles than any other particle sizes, a feature not observed in wild-type cells **(Figure 1 G-J)**.

**Figure 1:**
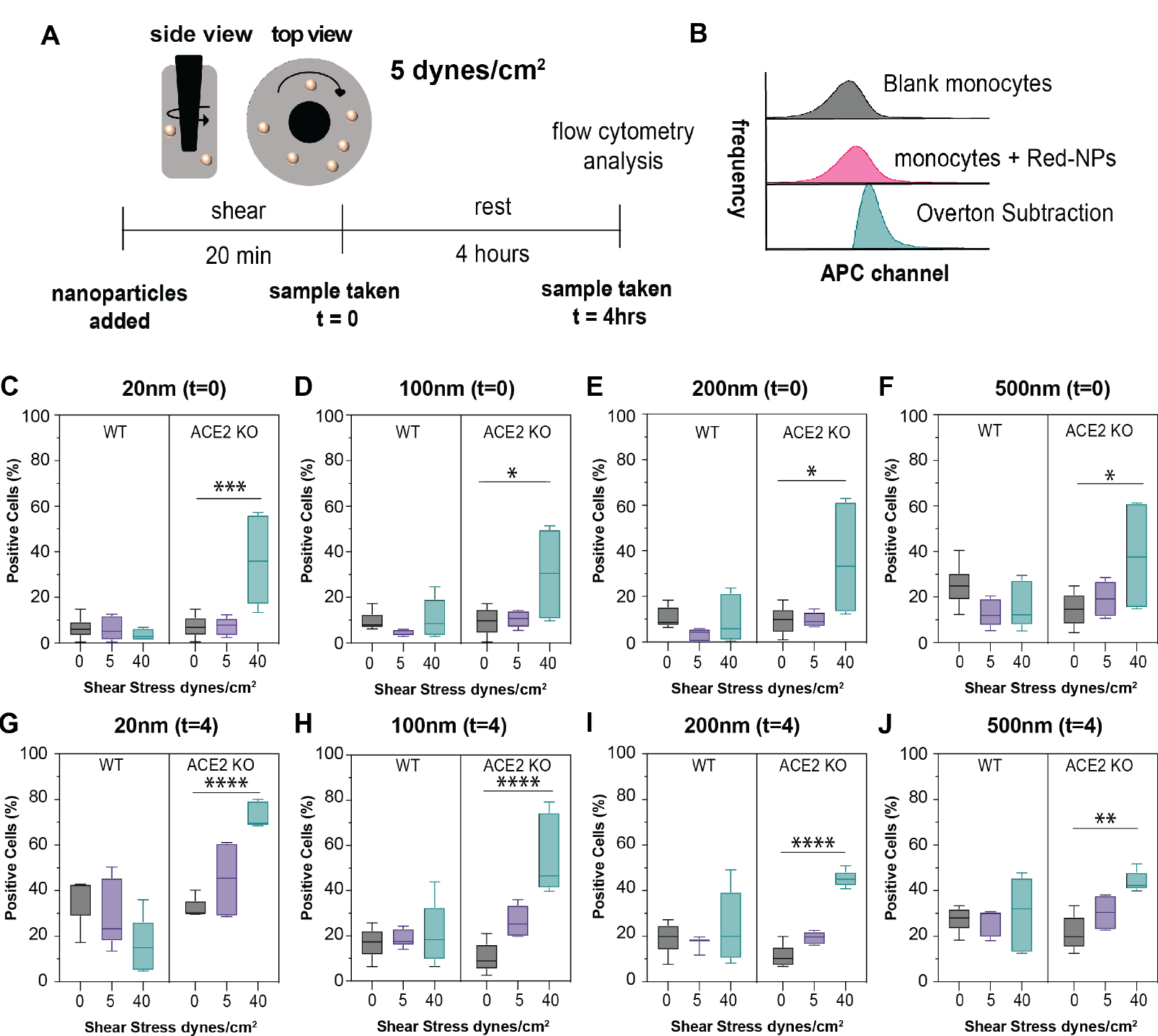
Percentage of wild type and ACE2-cells that took up particles during shear. a) schematic of rheometer geometry and timeline of shear experiments. (b) graphical visualization of flow cytometry quantifications. Subtraction histograms were used to determine the percentage of cells positive for nanoparticles compared to the control group of no particle and no shear. The percent positive cell of samples was measured directly after (t=0) the 20-minute exposure (C-F) or after a 4hr rest period (t=0) (G-J). Cells were exposed to no shear (static) (black), 5 dyne/cm^2^ (purple), or 40 dyne/cm^2^ (teal). Samples were taken in triplicate, and each replicate was added to the plot. Wild-type cells are a combination of 3 biological replicates, while ACE2^-^ cells are a combination of 2. Each graph represents one of the 4 sizes of nanoparticles tested at each time. Graphs are box and whisker plots, where the middle line represents the mean and error bars are the minimum and maximum of measured values. Statistical significance was determined using a 1-way ANOVA * P ≤ 0.05,** P ≤ 0.01, *** P ≤ 0.001 ^****^P ≤ 0.0001.

In wild-type cells, exposure to low or high shear did not significantly change the percentage of positive cells compared to static at any time point (Figure 1). Some diminishing was observed at 5 dyne/cm^2^ albeit not significantly. This difference increases with increasing size indicating the diminishing observed is exacerbated by larger particles (Figure 1).

In static and low shear conditions at t=0, no significant differences were observed between the percentage of nanoparticle-positive cells between WT and ACE2^-^ regardless of size (Figure 1). After the rest period (t=4), both cell types accumulated more particles but numbers between cell lines remained relatively similar indicating no difference in the sustained effect on the monocyte between the cell lines (Figure 1 G-J).

At higher shear, there is a significant difference in cell uptake between the two cell lines with 20 – 35% more ACE2^-^ cells taking up particles than WT at t=0. After the 4 hours of rest, the percentage of cells increased for each cell type, but the difference was relatively maintained with 20nm particles having the most difference increasing from approximately 30% to 50%.

Typically, classical monocytes are described as the endocytic phenotype^11^. The increase in ACE2^-^ cells which have taken up particles may be indicative of a more homogenous population of classical-like monocytes in higher shear regimes. This could also be an indication that lower shear in both cell types and high shear in WT cells induces more activation of monocytes skewing monocyte populations towards intermediate and non-classical phenotypes decreasing the number of cells in an endocytic phenotype. As patient plaque development progresses cells are going to experience higher shears due to the narrowing of the artery. Therefore, in a patient with a more progressed case of atherosclerosis, more cells may be interacting with nanotherapeutics. This could be of interest in therapeutic development because it could be useful when aiming for precision dosing of a patient.

Interestingly when cells were pre-treated with lipopolysaccharide (LPS) a comparable number of ACE2^-^ cells took up particles as WT cells directly after 40 dyne/cm^2^ of shear **(Supplementary Figure 1 A-D)**. This is contradictory to our finding under no pre-treatment conditions. LPS is an inflammatory stimulate skewing monocyte populations towards intermediate phenotypes. This could be the cause of the decreased number of cells taking up particles as cell phenotype skews from a phagocytic phenotype to more of a proliferative phenotype that drive an inflammatory response. Also of note is that at lower shears (5 dyne/cm2) a comparable number of cells took up more nanoparticles as cells exposed to high shear at t=0 (Supplementary Figure 2 A-D). This could be indicative of a similar inflammatory response from low shear and LPS stimulation.

After the rest period the comparable number of cells is diminished with ACE2^-^ cells having a larger number of cells positive for nanoparticles (Supplementary Figure 2 E-H). This may mean that the ACE2^-^ cells change phenotype more rapidly than WT cells.

### Wild type and ACE2^-^ cells have different genetic responses to shear stress despite similar uptake capacity

To better elucidate the impact of shear stress and nanoparticle size on monocyte endocytosis we measured the average mean intensity (MFIs) of a positive cell to determine the quantity of nanoparticles taken up per cell. Alongside these measurements, we tracked any phenotypic changes by measuring changes in monocyte genetic expression of adhesion molecules associated with Atherosclerosis, migration molecules typically associated with monocyte-endothelial interactions, and immune-modulating molecules indicative of monocyte phenotype **(Figures 2 & 3)**.

**Figure 2:**
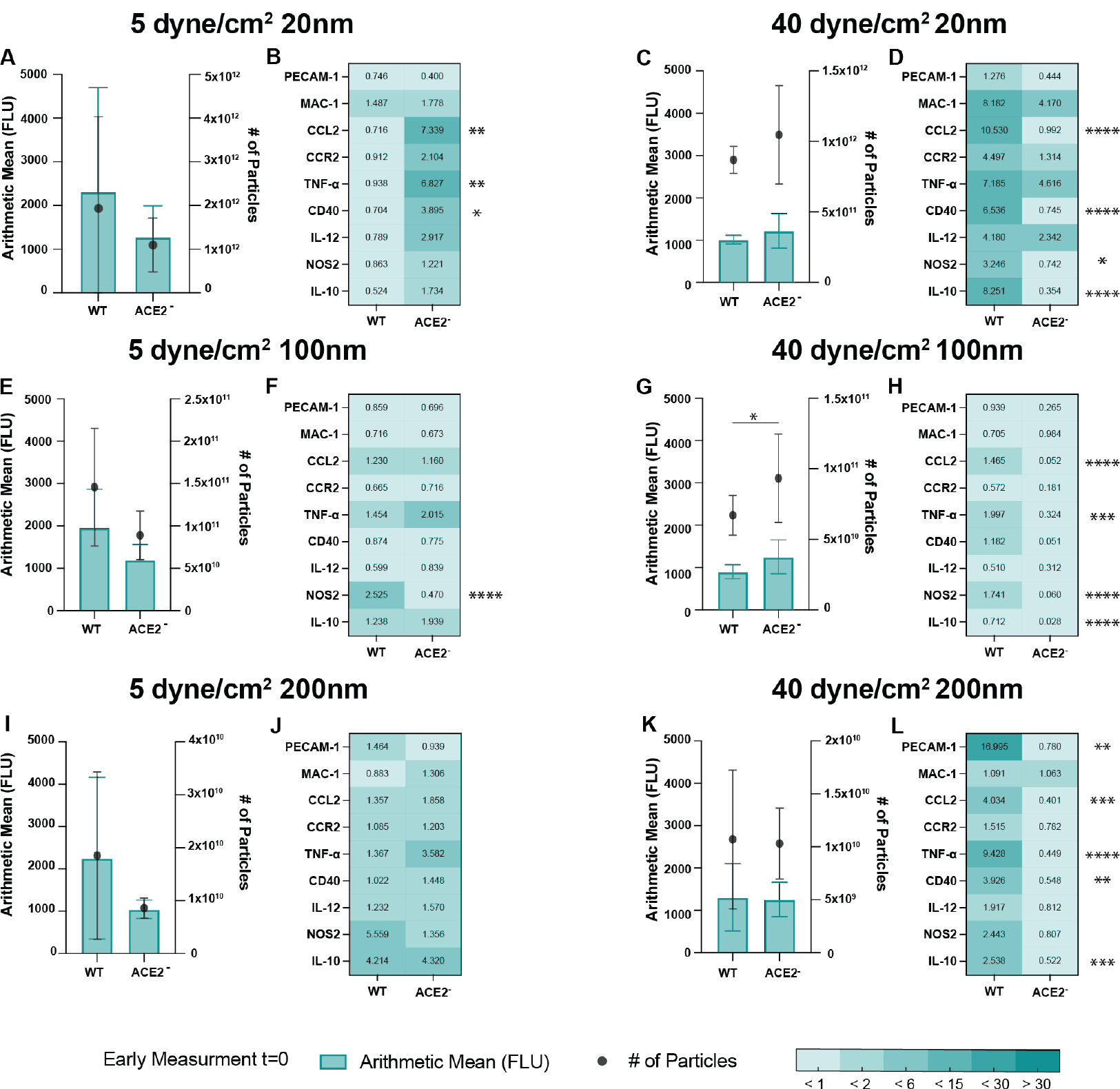
Average amount of particles taken up per cell and correlated genetic response directly after shear. Cells were exposed to 5 dyne/cm2 or 40 dyne/cm2 for 20 minutes. The mean fluorescence of the intensity per cell was calculated using the Overton method and is presented on the left Y axis (A, C, E, G, I, K). The number of particles/cells on the right Y axis was calculated using the calibration curve in Supplemental Figure 3 where fluorescent intensity was correlated with a known concentration of nanoparticles. Samples were taken in triplicate, and each replicate was added to the plot. Wild-type cells are a combination of 3 biological replicates, while ACE2^-^ cells are a combination of 2. Measurements were taken directly after the shear was applied (t=0). Statistical significance represented on the graph is between the number of particles of each sample. (B, D, F, H, J, L) Represent the genetic changes of the cells after the shear with nanoparticles. All values were calculated using the ΔΔCt method with the endogenous gene 18s and the reference sample was the respective cell line without nanoparticles and not exposed to shear. cDNA for qPCR was run in triplicate, and each value combines 2-3 biological replicates. Statistical significance was determined using a 2-way ANOVA * P ≤ 0.05, ** P ≤ 0.01,*** P ≤ 0.001, ^****^P ≤ 0.0001.

Across all sizes within the same cell type, we saw relatively similar mean fluorescence values, indicating that cells were taking up the same number of particles regardless of size. This is contradictory to most nanoparticle studies which have found that smaller nanoparticles are taken up more readily by monocytes^8^. We determined this was most likely due to the use of a consistent volume of 2% solid nanoparticle solution for all sizes instead of normalizing to particle number. To adjust a calibration curve was created which correlated fluorescent intensity to particle number **(Supplementary Figure 3)**.

Using this calibration curve, we could determine an estimate of how many particles each cell took up (Black circles in Figures 2 and 3). We found that 20nm particles were taken up in the largest quantity, followed by 100nm, and then 200nm as expected from previous literature.

**Figure 3:**
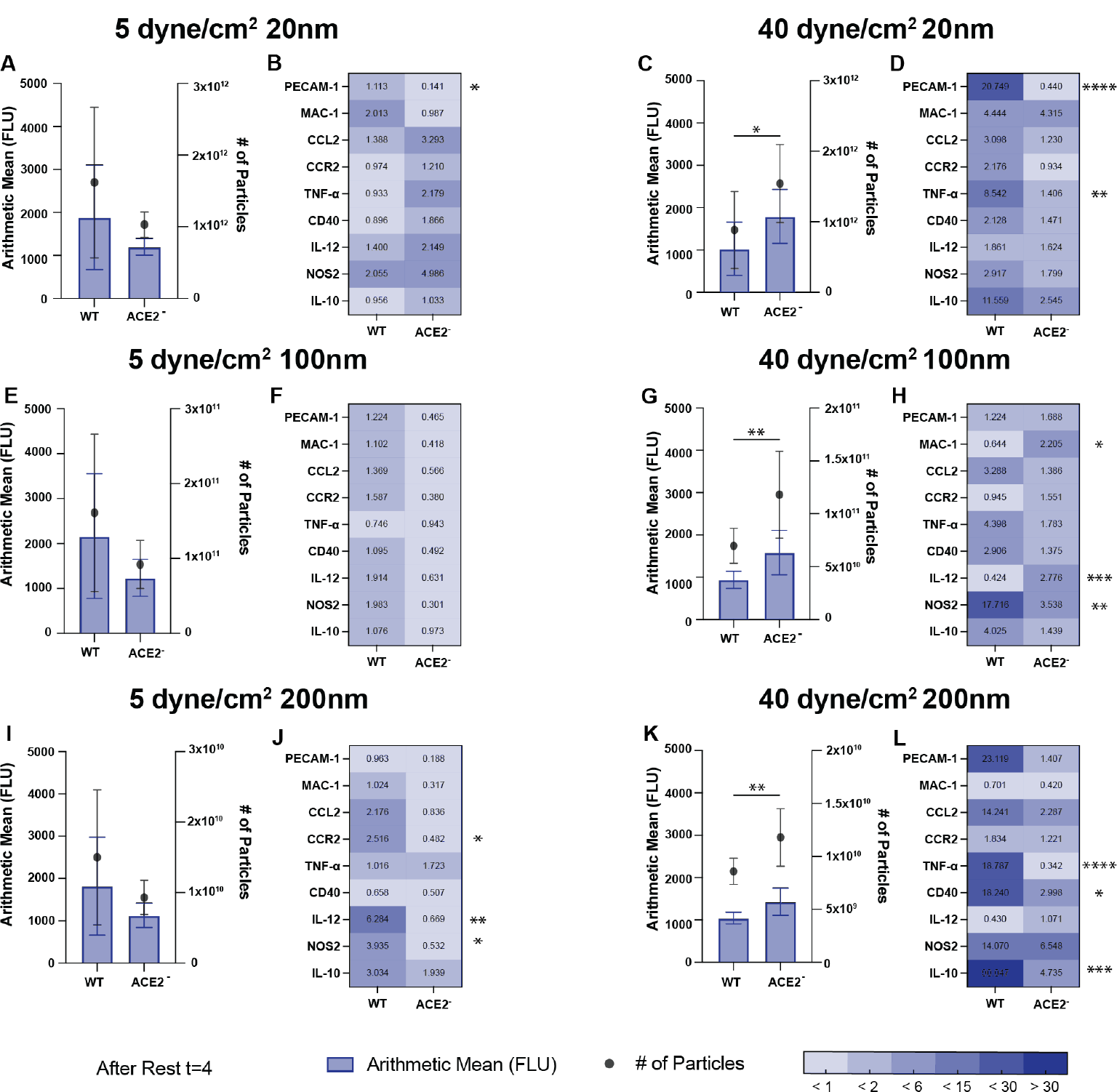
Average amount of particles taken up per cell and correlated genetic response after the 4-hour rest period. Cells were exposed to 5 dyne/cm2 or 40 dyne/cm2 for 20 minutes and then allowed to rest at 37^∘^C with 5% CO_2_. The mean fluorescence of the intensity per cell was calculated using the Overton method and is presented on the left Y axis (A, C, E, G, I, K). The number of particles/cells on the right Y axis was calculated using the calibration curve in Supplemental Figure 3 where fluorescent intensity was correlated with a known concentration of nanoparticles. Samples were taken in triplicate, and each replicate was added to the plot. Wild-type cells are a combination of 3 biological replicates, while ACE2^-^ cells are a combination of 2. Measurements were taken 4 hours after the shear was applied (t=4). Statistical significance represented on the graph is between the number of particles of each sample. (B, D, F, H, J, L) Represent the genetic changes of the cells after the shear with nanoparticles. All values were calculated using the ΔΔC_t_ method with the endogenous gene 18s and the reference sample was the respective cell line without nanoparticles and not exposed to shear. cDNA for qPCR was run in triplicate, and each value combines 2-3 biological replicates. Statistical significance was determined using a 2-way ANOVA * P ≤ 0.05, ** P ≤ 0.01, *** P ≤ 0.001, ^****^P ≤ 0.0001.

Similar to the results found in percent positive cells at lower shear, there are not many significant differences between the performance of WT cells and *ACE2*^*-*^ cells both initially after shear and after resting (Figure 2 A, E, I & Figure 3 A, E, I). Although not significant at the lower shear there is an observable decrease in the MFI of *ACE2*^*-*^ cells compared to WT in all sizes Figure 2 A, E, I & Figure 3 A, E, I). This indicates that both cell types take up a relatively similar number of particles. This is in good accordance with the findings in (Figure 1) where the same number of cells are also taking up cells. Genetic variation between the two cell lines at low shear shows upregulation in inflammatory cytokines in *ACE2*^*-*^ cells despite the lower MFI (Figure 2 B, F, J& Figure 3 B, F, J). Notably at t=0 with 20nm particles, there is significant upregulation of *CCL2, TNF-α*, and *CD40* when compared to WT which is lost at t=4 (Figure 2B & Figure 3B). Low shear with 100nm particles at t=0 did not have many significant differences in gene expression between cell lines but did show a significant upregulation of the *NOS2* in WT cells (Figure 2F) which indicates cells may have been taking on a phenotype that could protect against atherosclerosis by inhibiting adhesion and migration of leukocytes.^38^

At higher shear, both time points did not show any significant differences between the MFIs of WT and ACE2^-^ cells (Figure 2 C, G, K and Figure 3 C, G, K). When using the calibration curve to determine the number of particles per cell it was seen that at t=4 for all sizes ACE2^-^ cells have significantly more particles per cell but MFI differences remained insignificant (Figure 3 C, G, K). The lack of significant difference in MFI and particle number (apart from 100nm at t=0) is surprising due to the finding in (Figure 1) that shows that significantly more *ACE2*^*-*^ cells take up particles at higher shears. These findings together tell that a higher number of *ACE2*^*-*^ cells take up particles but the amount they take is comparable to WT cells. This further solidifies the hypothesis that *ACE2*^*-*^ cells develop a more homogenous population of endocytic monocytes under high shear than WT. At higher shear, there are more significantly different genes between the two cell types both at t=0 and t=4 than was found at lower shear (Figure 2 D, H, L & Figure 3 D, H, L). Upregulation of most genes was observed more in WT cells rather than *ACE2*^*-*^ cells (Figure 2 D, H, L & Figure 3 D, H, L). Much like 20nm at t=0 in low shear *CCL2* and *CD40* were among the upregulated genes in WT (Figure 2D). In contrast to the inflammatory nature of *CCL2* and *CD40*, there was also significant upregulation of *NOS2* and *IL-10* which generally perform more anti-inflammatory roles (Figure 2D). This trend of upregulation of both inflammatory and anti-inflammatory cytokines in WT cells at higher shear is carried through all sizes as well as at both time points (Figure 2 D, H, L & Figure 3 D, H, L).

An interesting trend that is emerging between the two cell types is the upregulation of genes in the cell line which have observably smaller MFI and number of particles. This along with the other findings indicates that cell phenotype has a higher impact on genetic response to shear than on its capacity to take up nanoparticles. It could also be an indication of altered phenotypic populations within an ACE2^-^ cell line when compared to WT.

Independent of nanoparticle exposure, we performed a proteomic analysis and found that shear stress induced changes in protein express **(Supplemental Figure 2)**. In comparison to wildtype THP1 monocytes, ACE2-THP1 monocytes showed increased expression of proteins related to migration, adhesion. In addition, proteins involved in ER stress response were also elevated at 40 dynes/cm^2^ compared to static and 5 dynes/cm^2^ conditions **(Supplemental Table 3)**. While these results are not directly related to endocytosis pathways, they are a good description of the different ways the cells respond to shear stress. These results highlight the complexity of shear stress, monocyte phenotype, and nanoparticle endocytosis on activation and uptake of particles.

### Using Augmented Design of Experiment Regressions to determine that shear stress and cell type are significant factors in nanoparticle endocytosis

Due to the large number of experimental conditions, we utilized statistical analysis software (SAS), JMP 17, to generate Standard Least Squares Linear Regression (SLSLR) models for four two-level, four-factor main effects screening design of experiments (12 runs each), and their corresponding DoEs. The four selected model parameters were the THP-1 cell type (wild-type or ACE2^-^), the shear stress (5 dyne/cm2 or 40 dyne/cm2) to which the monocytes were exposed, the size of the polystyrene nanoparticle (20, 100, 200, 500nm) in the culture, and the Resting Time (0 hr or 4 hr) before flow cytometric analysis. Each of the four main effects DOEs vary in the two levels selected for Nanoparticle Size: Model A (20 nm vs 100 nm), Model B (20 nm vs 200 nm), Model C (100 vs 200), and Model D (100 nm vs 500 nm). This allowed us to investigate monocytes’ uptake mechanism selectivity for macro-pinocytosis (20 nm), Clathrin-dependent endocytosis (100 nm), Caveolin receptor-mediated endocytosis (100 nm and 200 nm), or phagocytosis (500 nm).

Supplementary Table 4 summarizes the contribution of each parameter to the SLSLR models of each DOE (β coefficient estimates, P-values), the intercept (B0-estimate and Prob>|t|), and the overall model performance (R2, Adj-R2, RMSE, and P-Value). Models C and D are statistically significant, with P-values equal to 0.0332 and 0.0330, respectively; Model A has high statistical significance (P = 0.0022). Of the three statistically relevant models, Model D had the smallest root-mean-square error (RMSE) of 5.654, followed by Model A (RMSE=12.07), and Model C (RMSE=14.03; Table S1).

Due to the day-to-day variation of cellular performance in biological replicates, we wanted to improve upon the DOEs to encapsulate more of this variability. To improve these models, we implemented an augmented design to add two additional replicates of each of the original 12 runs (36 runs total/model). Descriptions of each run are provided Methods section. Table 1 summarizes the contribution of each parameter to the SLSLR models of each augmented DOE (β coefficient estimates, P-values), the intercept (B0-estimate and Prob>|t|), and the overall model performance (R2, Adj-R2, RMSE, and P-Value). Actual (experimentally obtained values) vs Predicted (data points based on linear regression model equation) plots are presented in Supplemental Figure 4.

**Table 1:**
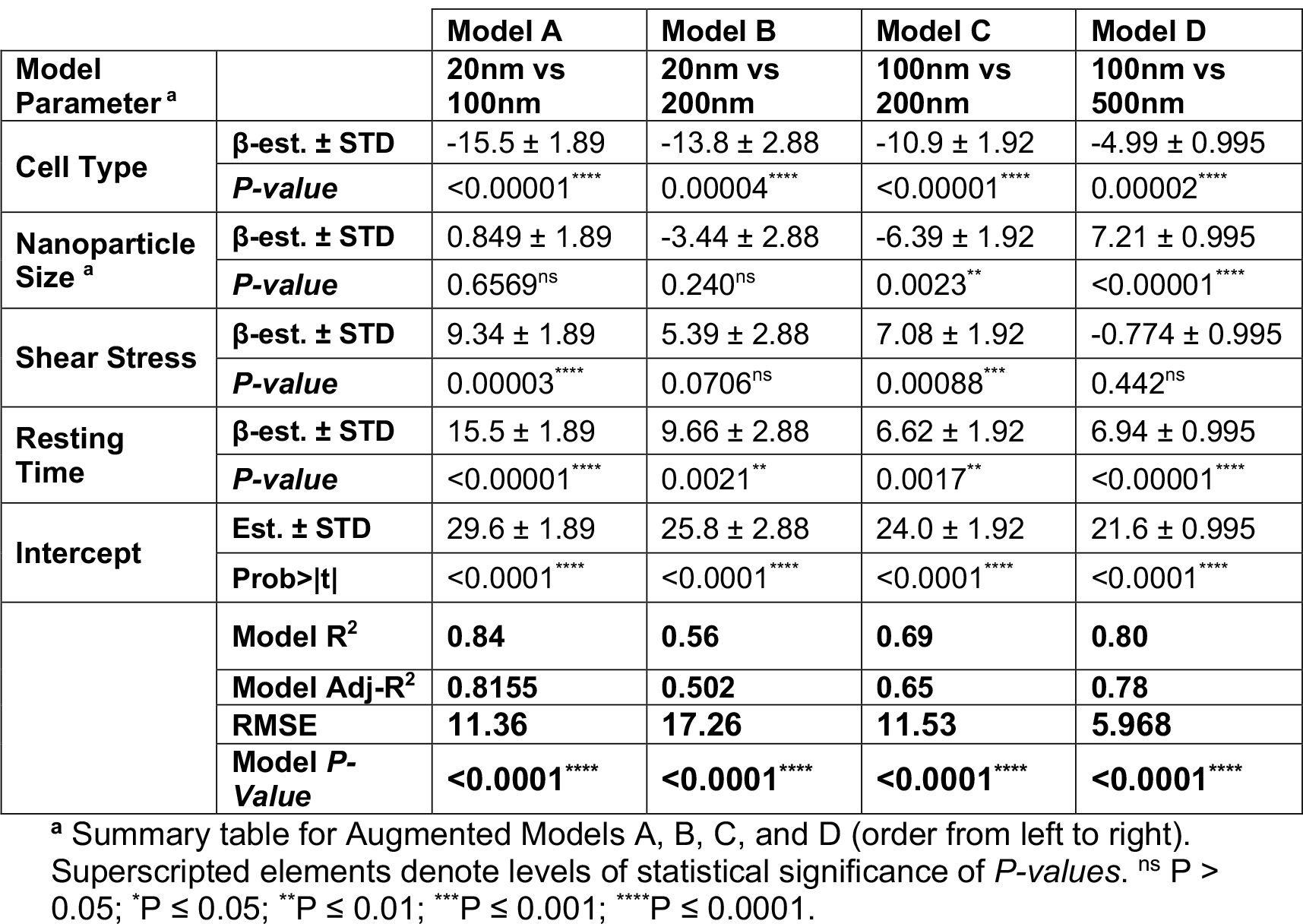
Values generated by JMP 17 SAS Standard Least Squares Linear Regression fit model DOE analysis. Overall model performance is described by correlation coefficients (R^2^), adjusted R^2^ (Adj-R^2^), root-mean-square error (RMSE), and *P-Values*.

All four Augmented Models had extremely significant Model P-Values (P < 0.0001 for all). Only one Augmented design, Model C, had a statistically significant contribution from each parameter to its linear regression model (Table 1). Matching results from the un-augmented design (Supplemental Table 4), Model D has the lowest RMSE of 5.968 (Table 1). Model A and D had one insignificant parameter (nanoparticle size and shear stress, respectively), while Model B had two insignificant parameters (also nanoparticle size and shear stress). Cell type and resting time factors contributed significantly to each of the four augmented designs. Interestingly, nanoparticle size is highly relevant (P = 0.0023) in Model C, where we are investigating Clathrin-vs Caveloin-mediated endocytosis. In this model, NP size has an estimated β-coefficient of -6.39 ± 1.92, indicating that nanoparticle size has an inverse correlation with the percent of cells positive for NP uptake (I.e., percent positive cells decrease as NP size increases, and vice versa). NP size is also extremely relevant (P > 0.00001) in Model D, where we investigate receptor-mediated endocytosis vs phagocytosis. In this model, NP size β-coefficient estimate is 7.21 ± 0.995. This suggests that the percent of positive cells increases with increasing NP size opposite of what we observe in Model C with Clathrin vs Caveloin endocytosis.

## Conclusion

Currently, there are no formal studies on how various nanoparticles activate monocytes, and many experiments vary their exposure conditions, making them difficult to compare 14. Moreover, studies also look at models with healthy monocytes, but there is information to be gained from understanding how activation and dynamics change in diseased systems 14. Our research highlights how studying multiple factors of a disease *in vitro* studies can change the expected outcomes from prior knowledge. This complex relationship between nanoparticle size, composition, shear, and monocyte phenotype is highlighted to show how short instances of shear can change monocyte behavior. This model while basic and hindered by the lack of ability of sustained long-term shear is the beginning of developing *in vitro* models which can be leveraged to make *in vitro* models which are more relevant to the development of disease-specific nanotherapeutics. With more robust *in vitro* models we can improve *in vivo* translation success. These more robust models can also be used to better understand monocyte phenotypic composition in diseases to better improve precision dosing for patients due to their specific needs.

## Acknowledgment

We would like to thank our funding sources National Institute of General Medical Sciences of the National Institutes of Health under award numbers 1 R35 GM142957-01, 5T32GM133353-02, and the GEM Fellowship. Also a special thanks to Hannah Yankello, and Connor Valentine for support in learning the Flow cytometry and Rheology Techniques respectively.

---

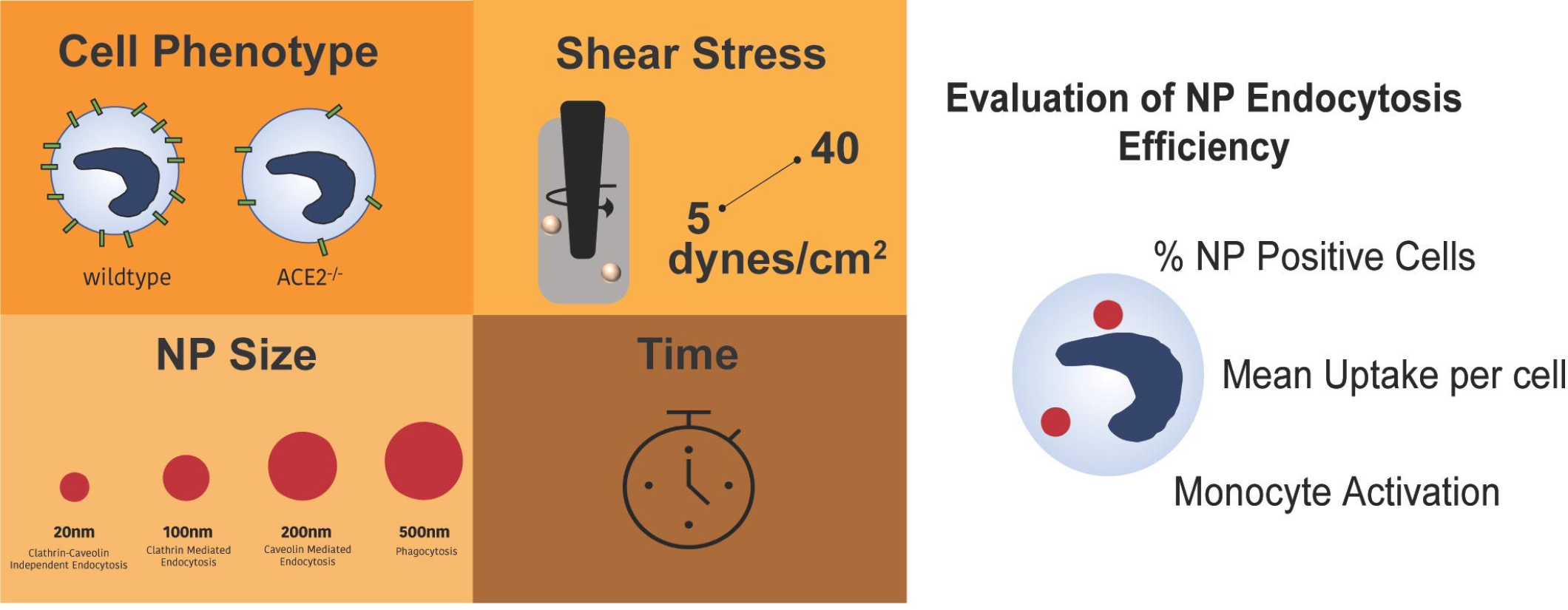

## Notes

### Competing Interest Statement

The authors have declared no competing interest.

