## Supplemental Figures for "Nanoparticle endocytosis is driven by monocyte phenotype rather than nanoparticle size under high shear flow conditions"

**Supplementary Table 1: Raw data used for 36-run Augmented Design Models A and B**

| | Model A | | | | | Model B | | | | |
|---|---|---|---|---|---|---|---|---|---|---|
| Run # | Shear Stress | NP Size | Cell type | Resting Time | Percent positive cells | Shear Stress | NP Size | Cell Type | Resting Time | Percent positive cells |
| 1 | 5 | 20 | wt | 0 | 1.39 | 5 | 100 | wt | 4 | 23.04 |
| 2 | 5 | 20 | wt | 0 | 5.18 | 5 | 100 | wt | 4 | 24.26 |
| 3 | 5 | 20 | wt | 0 | 3.37 | 5 | 100 | wt | 4 | 23.23 |
| 4 | 40 | 20 | wt | 0 | 1.27 | 40 | 100 | wt | 4 | 6.34 |
| 5 | 40 | 20 | wt | 0 | 1.36 | 40 | 100 | wt | 4 | 11.13 |
| 6 | 40 | 20 | wt | 0 | 0 | 40 | 100 | wt | 4 | 10.74 |
| 7 | 40 | 100 | wt | 0 | 2.92 | 40 | 100 | wt | 0 | 3.45 |
| 8 | 40 | 100 | wt | 0 | 11.74 | 40 | 100 | wt | 0 | 17.26 |
| 9 | 40 | 100 | wt | 0 | 24.74 | 40 | 100 | wt | 0 | 5.32 |
| 10 | 5 | 20 | ACE2 | 0 | 2.46 | 5 | 500 | wt | 4 | 30.74 |
| 11 | 5 | 20 | ACE2 | 0 | 3.52 | 5 | 500 | wt | 4 | 30.52 |
| 12 | 5 | 20 | ACE2 | 0 | 5.74 | 5 | 500 | wt | 4 | 30.67 |
| 13 | 5 | 100 | ACE2 | 0 | 14.28 | 5 | 500 | wt | 0 | 20.49 |
| 14 | 5 | 100 | ACE2 | 0 | 13.83 | 5 | 500 | wt | 0 | 18.39 |
| 15 | 5 | 100 | ACE2 | 0 | 13.55 | 5 | 500 | wt | 0 | 20.24 |
| 16 | 40 | 100 | ACE2 | 0 | 49 | 40 | 500 | wt | 0 | 8.68 |
| 17 | 40 | 100 | ACE2 | 0 | 48.23 | 40 | 500 | wt | 0 | 5.23 |
| 18 | 40 | 100 | ACE2 | 0 | 51.34 | 40 | 500 | wt | 0 | 9.96 |
| 19 | 40 | 20 | wt | 4 | 23.11 | 5 | 100 | ACE2 | 4 | 19.87 |
| 20 | 40 | 20 | wt | 4 | 22.9 | 5 | 100 | ACE2 | 4 | 19.83 |
| 21 | 40 | 20 | wt | 4 | 35.94 | 5 | 100 | ACE2 | 4 | 20.05 |
| 22 | 5 | 100 | wt | 4 | 23.04 | 5 | 100 | ACE2 | 0 | 14.28 |
| 23 | 5 | 100 | wt | 4 | 24.26 | 5 | 100 | ACE2 | 0 | 13.83 |
| 24 | 5 | 100 | wt | 4 | 23.23 | 5 | 100 | ACE2 | 0 | 13.55 |
| 25 | 5 | 100 | wt | 4 | 17.51 | 40 | 100 | ACE2 | 0 | 10.87 |
| 26 | 5 | 100 | wt | 4 | 16.7 | 40 | 100 | ACE2 | 0 | 9.83 |
| 27 | 5 | 100 | wt | 4 | 14.11 | 40 | 100 | ACE2 | 0 | 12.81 |
| 28 | 5 | 20 | ACE2 | 4 | 61.05 | 5 | 500 | ACE2 | 0 | 26.44 |
| 29 | 5 | 20 | ACE2 | 4 | 60.54 | 5 | 500 | ACE2 | 0 | 25.55 |
| 30 | 5 | 20 | ACE2 | 4 | 60.59 | 5 | 500 | ACE2 | 0 | 28.5 |
| 31 | 40 | 20 | ACE2 | 4 | 79.02 | 40 | 500 | ACE2 | 4 | 41.17 |
| 32 | 40 | 20 | ACE2 | 4 | 69.66 | 40 | 500 | ACE2 | 4 | 42.53 |
| 33 | 40 | 20 | ACE2 | 4 | 80.11 | 40 | 500 | ACE2 | 4 | 41.96 |
| 34 | 40 | 100 | ACE2 | 4 | 73.03 | 40 | 500 | ACE2 | 4 | 46.83 |
| 35 | 40 | 100 | ACE2 | 4 | 79.24 | 40 | 500 | ACE2 | 4 | 39.81 |
| 36 | 40 | 100 | ACE2 | 4 | 47.03 | 40 | 500 | ACE2 | 4 | 51.69 |

**Supplementary Table 2: Raw data used for 36-run Augmented Design Models C and D**

| | Model C | | | | | Model D | | | | |
|---|---|---|---|---|---|---|---|---|---|---|
| Run # | Shear Stress | NP Size | Cell type | Resting Time | Percent positive cells | Shear Stress | NP Size | Cell Type | Resting Time | Percent positive cells |
| 1 | 5 | 100 | ACE2 | 0 | 14.28 | 5 | 20 | wt | 0 | 1.39 |
| 2 | 5 | 100 | ACE2 | 0 | 13.83 | 5 | 20 | wt | 0 | 5.18 |
| 3 | 5 | 100 | ACE2 | 0 | 13.55 | 5 | 20 | wt | 0 | 3.37 |
| 4 | 5 | 100 | wt | 4 | 23.04 | 40 | 20 | wt | 0 | 1.36 |
| 5 | 5 | 100 | wt | 4 | 24.26 | 40 | 20 | wt | 0 | 1.27 |
| 6 | 5 | 100 | wt | 4 | 23.23 | 40 | 20 | wt | 0 | 0 |
| 7 | 5 | 100 | wt | 4 | 17.51 | 5 | 200 | wt | 0 | 14.38 |
| 8 | 5 | 100 | wt | 4 | 16.7 | 5 | 200 | wt | 0 | 13.5 |
| 9 | 5 | 100 | wt | 4 | 14.11 | 5 | 200 | wt | 0 | 13.66 |
| 10 | 40 | 100 | wt | 0 | 24.74 | 5 | 20 | ACE2 | 0 | 6.72 |
| 11 | 40 | 100 | wt | 0 | 2.92 | 5 | 20 | ACE2 | 0 | 6.52 |
| 12 | 40 | 100 | wt | 0 | 11.74 | 5 | 20 | ACE2 | 0 | 6.88 |
| 13 | 40 | 100 | ACE2 | 0 | 51.34 | 40 | 200 | ACE2 | 0 | 13.46 |
| 14 | 40 | 100 | ACE2 | 0 | 49 | 40 | 200 | ACE2 | 0 | 12.17 |
| 15 | 40 | 100 | ACE2 | 0 | 48.23 | 40 | 200 | ACE2 | 0 | 18.82 |
| 16 | 40 | 100 | ACE2 | 4 | 73.03 | 40 | 200 | ACE2 | 0 | 47.52 |
| 17 | 40 | 100 | ACE2 | 4 | 79.24 | 40 | 200 | ACE2 | 0 | 60.68 |
| 18 | 40 | 100 | ACE2 | 4 | 47.03 | 40 | 200 | ACE2 | 0 | 62.88 |
| 19 | 5 | 200 | wt | 0 | 14.38 | 40 | 20 | wt | 4 | 23.11 |
| 20 | 5 | 200 | wt | 0 | 13.5 | 40 | 20 | wt | 4 | 22.9 |
| 21 | 5 | 200 | wt | 0 | 13.66 | 40 | 20 | wt | 4 | 35.94 |
| 22 | 5 | 200 | ACE2 | 0 | 12.48 | 5 | 200 | wt | 4 | 17.7 |
| 23 | 5 | 200 | ACE2 | 0 | 10.87 | 5 | 200 | wt | 4 | 17.97 |
| 24 | 5 | 200 | ACE2 | 0 | 14.46 | 5 | 200 | wt | 4 | 11.68 |
| 25 | 5 | 200 | ACE2 | 4 | 21.62 | 40 | 200 | wt | 4 | 12.37 |
| 26 | 5 | 200 | ACE2 | 4 | 21.33 | 40 | 200 | wt | 4 | 8.22 |

| | | | | | | | | | | |
|---|---|---|---|---|---|---|---|---|---|---|
| 27 | 5 | 200 | ACE 2 | 4 | 22.58 | 40 | 200 | wt | 4 | 11.08 |
| 28 | 40 | 200 | wt | 0 | 1.02 | 5 | 20 | ACE2 | 4 | 61.05 |
| 29 | 40 | 200 | wt | 0 | 3.33 | 5 | 20 | ACE2 | 4 | 60.54 |
| 30 | 40 | 200 | wt | 0 | 0.34 | 5 | 20 | ACE2 | 4 | 60.59 |
| 31 | 40 | 200 | wt | 4 | 12.37 | 40 | 20 | ACE2 | 4 | 79.02 |
| 32 | 40 | 200 | wt | 4 | 8.22 | 40 | 20 | ACE2 | 4 | 69.66 |
| 33 | 40 | 200 | wt | 4 | 11.08 | 40 | 20 | ACE2 | 4 | 80.11 |
| 34 | 40 | 200 | ACE 2 | 4 | 44.68 | 5 | 200 | ACE2 | 4 | 21.62 |
| 35 | 40 | 200 | ACE 2 | 4 | 40.87 | 5 | 200 | ACE2 | 4 | 21.33 |
| 36 | 40 | 200 | ACE 2 | 4 | 50.94 | 5 | 200 | ACE2 | 4 | 22.58 |

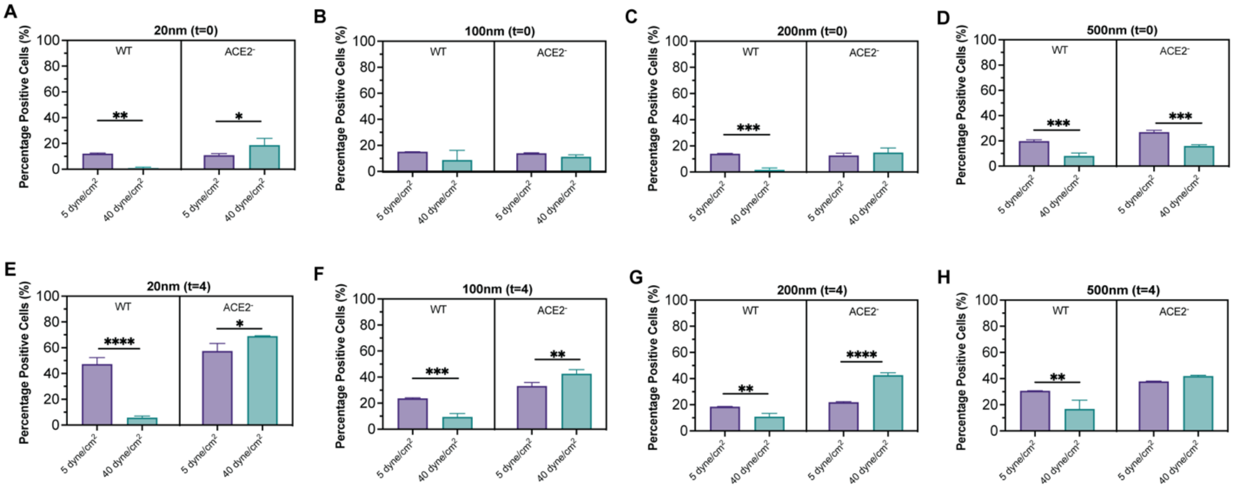


**Supplemental Figure 1: Percentage of LPS Stimulated THP-1 Cells that took up nanoparticles under shear.** Subtraction histograms were used to determine the percentage of cells positive for nanoparticles compared to the control group of no particles and no shear. The percent positive cell of samples was measured directly after (t=0) the 20-minute exposure (C-F) or after a 4hr rest period (t=4) (E-H). Cells were exposed to either 5 dyne/cm$^2$ (purple) or 40 dyne/cm$^2$ (teal). Samples were taken in triplicate, and each replicate was added to the plot. Each graph represents one of the 4 sizes of nanoparticles tested at each time. The graphs are box and whisker plots, where the middle line represents the mean, and the error bars are the minimum and maximum of measured values. Statistical significance was determined using a 1-way ANOVA * P ≤ 0.05, ** P ≤ 0.01, *** P ≤ 0.001, **** P ≤ 0.0001.

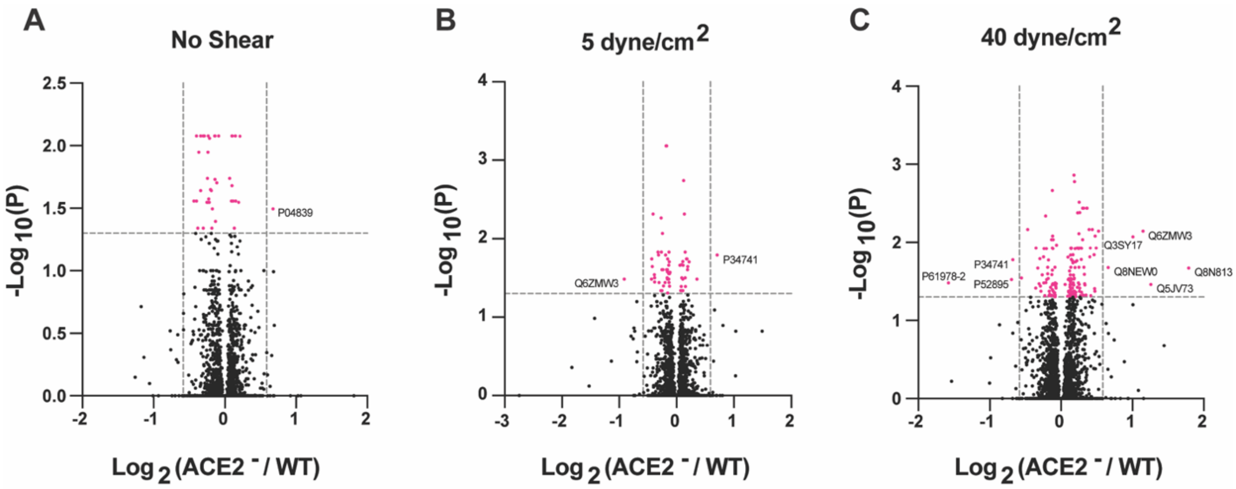


**Supplemental Figure 2: Differential protein expression analysis between THP1 wildtype and ACE2- monocytes under physiological shear stress.** THP1 wildtype cells were compared to ACE2 deficient monocytes in their ability to endocytose nanoparticles prior to any shear applied (A), or after dynes/cm$^2$ (B) or 40 dynes/cm$^2$ (C). Following the shear experiments, cell lysate was isolated and prepared for proteomic analysis. Volcano plots to identify significantly regulated proteins (fold change equal to or greater than ∓ 1.5, P ≤ 0.05). All comparisons use wt are the reference with N=3 technical replicates per sample group.

| | | Shear Stress (dyne/cm$^2$) | | |
|---|---|---|---|---|
| **Accession** | **Name** | **0** | **5** | **40** |
| P04839 | Cytochrome b-245 heavy chain | Up | -- | -- |
| Q6ZMW3 | Echinoderm microtubule-associated protein-like 6 | -- | Down | -- |
| P34741 | Syndecan-2 | -- | Up | Down |
| P52895 | Aldo-keto reductase family 1 member C2 | -- | -- | Down |
| P61978-2 | Heterogeneous nuclear ribonucleoprotein K | -- | -- | Down |
| Q3SY17 | Mitochondrial nicotinamide adenine dinucleotide transporter SLC25A52 | -- | -- | Up |
| Q8NEW0 | Zinc transporter 7 | -- | -- | Up |
| Q6ZMW3 | Echinoderm microtubule-associated protein-like 6 | -- | -- | Up |
| Q5JV73 | FERM and PDZ domain-containing protein 3 | -- | -- | Up |
| Q8N813 | Putative uncharacterized protein C3orf56 | -- | -- | Up |

**Supplementary Table 3: Most significantly changed proteins between THP1 wildtype and *ACE2*- monocytes.** Under physiological shear stress. – denotes no change. “Up” and “Down” are all relative to *ACE2*- monocytes.

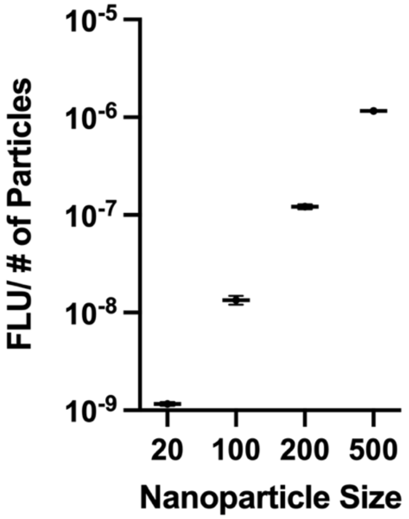


**Supplemental Figure 3: Fluorescence per number of particle calibration curve.** Samples of known concentrations of nanoparticles were measured on a fluorescent plate reader. Fluorescence was then divided by the known number of particles. Error bars represent the variation of samples from a dilution series of the original sample.

| | | Model A | Model B | Model C | Model D |
|---|---|---|---|---|---|
| **Model Parameter** | | **20nm vs. 100nm** | **20nm vs. 200nm** | **100nm vs. 200nm** | **100nm vs. 500nm** |
| **Cell Type** | **β-est. ± STD** | -15.5 ± 3.48 | -13.1 ± 6.36 | -11.4 ± 4.05 | -5.20 ± 1.63 |
| | ***P-value*** | 0.0030$^{**}$ | 0.078$^{ns}$ | 0.026$^{*}$ | 0.015$^{*}$ |
| **Nanoparticle Size** | **β-est. ± STD** | 1.60 ± 3.48 | -3.84 ± 6.36 | -8.26 ± 4.05 | 7.46 ± 1.63 |
| | ***P-value*** | 0.659$^{ns}$ | 0.565$^{ns}$ | 0.081$^{ns}$ | 0.0026$^{**}$ |
| **Shear Stress** | **β-est. ± STD** | 12.4 ± 3.48 | 6.58 ± 6.36 | 9.22 ± 4.05 | -1.88 ± 1.63 |
| | ***P-value*** | 0.0093$^{**}$ | 0.335$^{ns}$ | 0.057$^{ns}$ | 0.286$^{ns}$ |
| **Resting Time** | **β-est. ± STD** | 15.8 ± 3.48 | 10.3 ± 6.36 | 6.31 ± 4.05 | 6.47 ± 1.63 |
| | ***P-value*** | 0.0027$^{**}$ | 0.150$^{ns}$ | 0.163$^{ns}$ | 0.0054$^{**}$ |
| **Intercept** | **Est. ± STD** | 32.9 ± 3.48 | 27.6 ± 6.36 | 26.2 ± 4.05 | 20.6 ± 1.63 |
| | **Prob>\|t\|** | <0.0001**** | <0.0001**** | <0.0001**** | <0.0001**** |
| | **Model $R^2$** | **0.88** | **0.54** | **0.74** | **0.87** |
| | **Model Adj-$R^2$** | **0.82** | **0.28** | **0.59** | **0.80** |
| | **RMSE** | **12.07** | **22.03** | **14.03** | **5.654** |
| | **Model *P-Value*** | **0.0022$^{**}$** | **0.1878$^{ns}$** | **0.0332$^{*}$** | **0.0330*** |

**Supplementary Table 4: Values generated by JMP 17 SAS Standard Least Squares Linear Regression fit model DOE analysis for un-augmented runs.** Overall model performance is described by correlation coefficients ($R^2$), adjusted $R^2$ (Adj-$R^2$), root-mean-square error (RMSE), and *P-Values*. Superscripted elements denote levels of statistical significance of *P-values*. $^{ns}$ P > 0.05; $^{*}$ P ≤ 0.05; $^{**}$ P ≤ 0.01; $^{***}$ P ≤ 0.001; $^{****}$ P ≤ 0.0001.

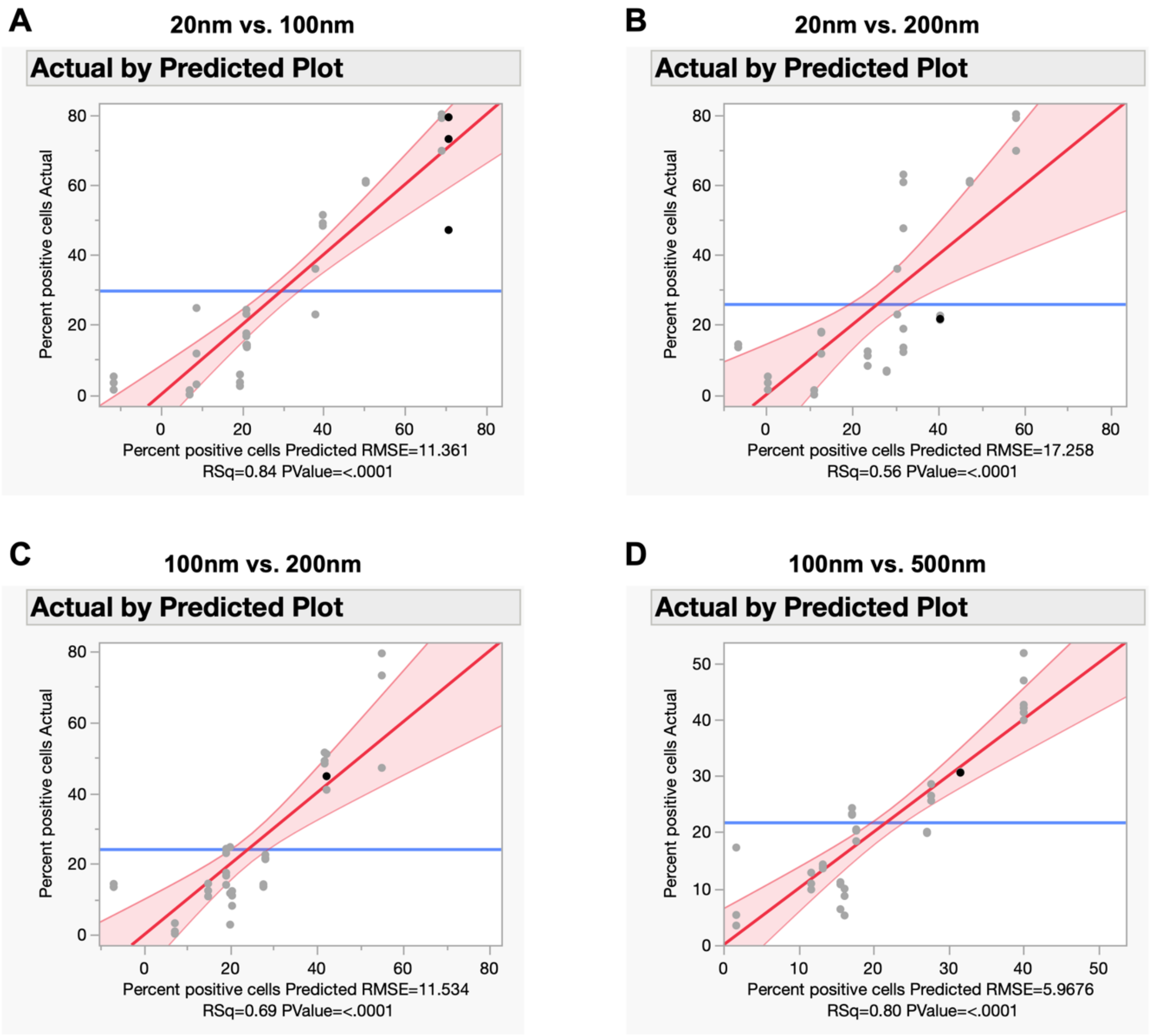


**Supplemental Figure 4: Actual vs predicted plots of the standard least squares regression models for percentage of THP-1 cells positive for nanoparticle uptake.** Each panel contains a single plot representing one of four Augmented DoEs, with each varying in the two levels of polystyrene NP diameters available for THP-1 monocyte uptake: (a) 20 nm vs 100 nm, (b) 20 nm vs 200 nm, (c) 100 nm vs 200 nm, and (d) 100 nm vs 500 nm.
